# Rev-erb-α controls skeletal muscle calcium homeostasis through myoregulin repression: implications in Duchenne Muscular Dystrophy

**DOI:** 10.1101/2021.04.22.440922

**Authors:** Alexis Boulinguiez, Christian Duhem, Alicia Mayeuf-Louchart, Benoit Pourcet, Yasmine Sebti, Kateryna Kondratska, Valérie Montel, Stéphane Delhaye, Quentin Thorel, Justine Beauchamp, Aurore Hebras, Marion Gimenez, Marie Couvelaere, Mathilde Zecchin, Lise Ferri, Natalia Prevarskaya, Anne Forand, Christel Gentil, France Piétri-Rouxel, Bruno Bastide, Bart Staels, Helene Duez, Steve Lancel

## Abstract

The sarcoplasmic reticulum (SR) plays an important role in calcium homeostasis. SR calcium mishandling is described in pathological conditions such as myopathies. Here, we investigated whether the nuclear receptor Rev-erb-α regulates skeletal muscle SR calcium homeostasis. Our data demonstrate that Rev-erbα invalidation in mice impairs SERCA-dependent SR calcium uptake. Rev-erb-α acts on calcium homeostasis by repressing the SERCA inhibitor Myoregulin, through direct binding to its promoter. Restoration of Myoregulin counteracts the effects of REV-ERB-α overexpression on SR calcium content. Interestingly, myoblasts from Duchenne myopathy patients display downregulated REV-ERBα expression, whereas pharmacological Rev-erb activation ameliorates SR calcium homeostasis, and improves muscle structure and function in dystrophic *mdx/Utr^+/-^* mice. Our findings demonstrate that Rev-erb-α regulates muscle SR calcium homeostasis, pointing to its therapeutic interest for mitigating myopathy.

## Introduction

Skeletal muscle is not only required for movements, but is also crucial for other vital functions such as respiration. Myopathies (Lee and Noguchi, 2016; Vallejo-Illarramendi et al., 2014), among which Duchenne Muscular Dystrophy (DMD) is one of the most prevalent forms, result in progressive muscle weakness and wasting, and can lead to premature death. Despite progress in gene therapy, DMD still remains an unmet medical need, calling for new strategies to alleviate skeletal muscle degeneration.

Calcium (Ca^2+^) is important for muscle contractile function and its subcellular distribution is tightly regulated by several pumps and channels (Calderón et al., 2014). Ca^2+^ is stored in the endoplasmic/sarcoplasmic reticulum (ER/SR) where it mainly interacts with Ca^2+^-binding proteins such as Calsequestrin (Michalak et al., 2009). Following an action potential, membrane depolarization triggers a massive Ca^2+^ release through the Ryanodine Receptor (RyR), hence promoting contraction and muscle force generation. Ca^2+^ reuptake from the cytosol into the SR lumen by the Sarco/Endoplasmic Reticulum Calcium ATPase (SERCA) allows muscle relaxation and a new cycle of contraction/relaxation. Because SERCA Ca^2+^ pump activity plays a prominent role in skeletal muscle contractility, it is tightly regulated by different factors including the recently discovered inhibitory micropeptide Myoregulin (Mln) (Anderson et al., 2015). Disturbances of these fine-tuned processes have been observed in DMD, where chronically elevated cytosolic Ca^2+^ concentrations (Farini et al., 2016; Mázala et al., 2015), decreased SERCA activity (Gehrig et al., 2012; Kargacin and Kargacin, 1996; Schneider et al., 2013) and reduced Ca^2+^ release upon excitation can be observed (Hollingworth et al., 2008; Kargacin and Kargacin, 1996; Vallejo-Illarramendi et al., 2014). Progressive loss of muscle force generation, as observed in the *mdx* mouse model of DMD, is explained by the absence of homeostatic return to basal cytosolic Ca^2+^ levels between two contractions (Claflin and Brooks, 2008), underlying the importance of normal SERCA activity for muscle function.

We have previously reported that the druggable nuclear receptor and transcriptional repressor Rev-erb-α (Harding and Lazar, 1995) improves skeletal muscle function and exercise capacity (Woldt et al., 2013). Especially, Rev-erb-α improves mitochondrial function along with increased mitochondrial biogenesis, and reduces autophagy (Woldt et al., 2013). We investigated here whether Rev-erb-α controls additional mechanisms accounting for skeletal homeostasis. We particularly assessed whether Rev-erb-α modulates the major SR function, *i.e.* Ca^2+^ handling. We demonstrate that Rev-erb-α, through its transcriptional repressive activity on the *Mln* gene, increases SERCA activity and SR Ca^2+^ content. Importantly, pharmacological Rev-erb activation with SR9009 decreases *Mln* expression, improves calcium handling, enhances force generation and minimizes tissue damage in severely dystrophic *mdx/utr^+/-^* mice. Overall, our results identify Rev-erb-α as a new regulator of SR calcium homeostasis that may represent a therapeutic target in skeletal muscle disorders related to impaired reticular calcium homeostasis, such as myopathies.

## Results

### Rev-erb-α improves muscle force along with better SR Ca^2+^ homeostasis

We first aimed to determine whether Rev-erb-a is important for muscle force generation and found that muscle contraction is reduced by ~50% in *Rev-erbα^-/-^* mice compared to their wild-type (*Rev-erbα*^+/+^) littermate controls (Figure 1A). Since Ca^2+^ homeostasis is essential for muscle force generation we next determined whether Rev-erb-α controls SR Ca^2+^ handling. Muscle microsomes, *i.e.* sarcoplasmic vesicles, were prepared from *Rev-erbα^-/-^* and *Rev-erbα^+/+^* littermates. SERCA-dependent SR Ca^2+^ uptake capacity was measured over time after the addition of Ca^2+^ pulses or thapsigargin (TG), a potent inhibitor of SERCA activity (Lytton et al., 1991), by using a fluorescent probe detecting extramicrosomal Ca^2+^. The slope of fluorescence decrease, *i.e.* SR Ca^2+^ uptake, was significantly lower in *Rev-erbα^-/-^* mice compared to *Rev-erbα^+/+^* mice, revealing a reduction in SERCA activity in absence of Rev-erb-α (Figures 1B and 1C). By contrast, muscle microsomes prepared from mice treated with the Rev-erb agonist SR9009 elicited improved skeletal muscle SERCA activity (Figures 1D and 1E). In order to measure passive Ca^2+^ release from SR as a surrogate of its initial Ca^2+^ content, differentiated *REV-ERBα*-overexpressing or control (pBabe) C2C12 myotubes were loaded with the cytosolic Ca^2+^-sensitive probe Fluo4-AM and then challenged with TG to release Ca^2+^ from the SR. TG addition led to a greater elevation in Fluo4 fluorescence in *REV-ERBα*-overexpressing cells compared to pBabe, revealing that Rev-erb-α overexpression is associated with increased SR Ca^2+^ content (Figures 1F and 1G). Consistently, similar results were obtained using the cytosolic Ca^2+^-sensitive probe Fura-2 AM, which is a dual-excitation, single-emission Ca^2+^ indicator avoiding possible loading artifacts (Figures 1H and 1I). Basal cytosolic calcium, buffered at least in part by the SR, is reduced by *REV-ERBα*-overexpression (Figure 1J). Together, these data indicate that Rev-erb-α controls SR Ca^2+^ homeostasis in skeletal muscle.

**Figure 1.**
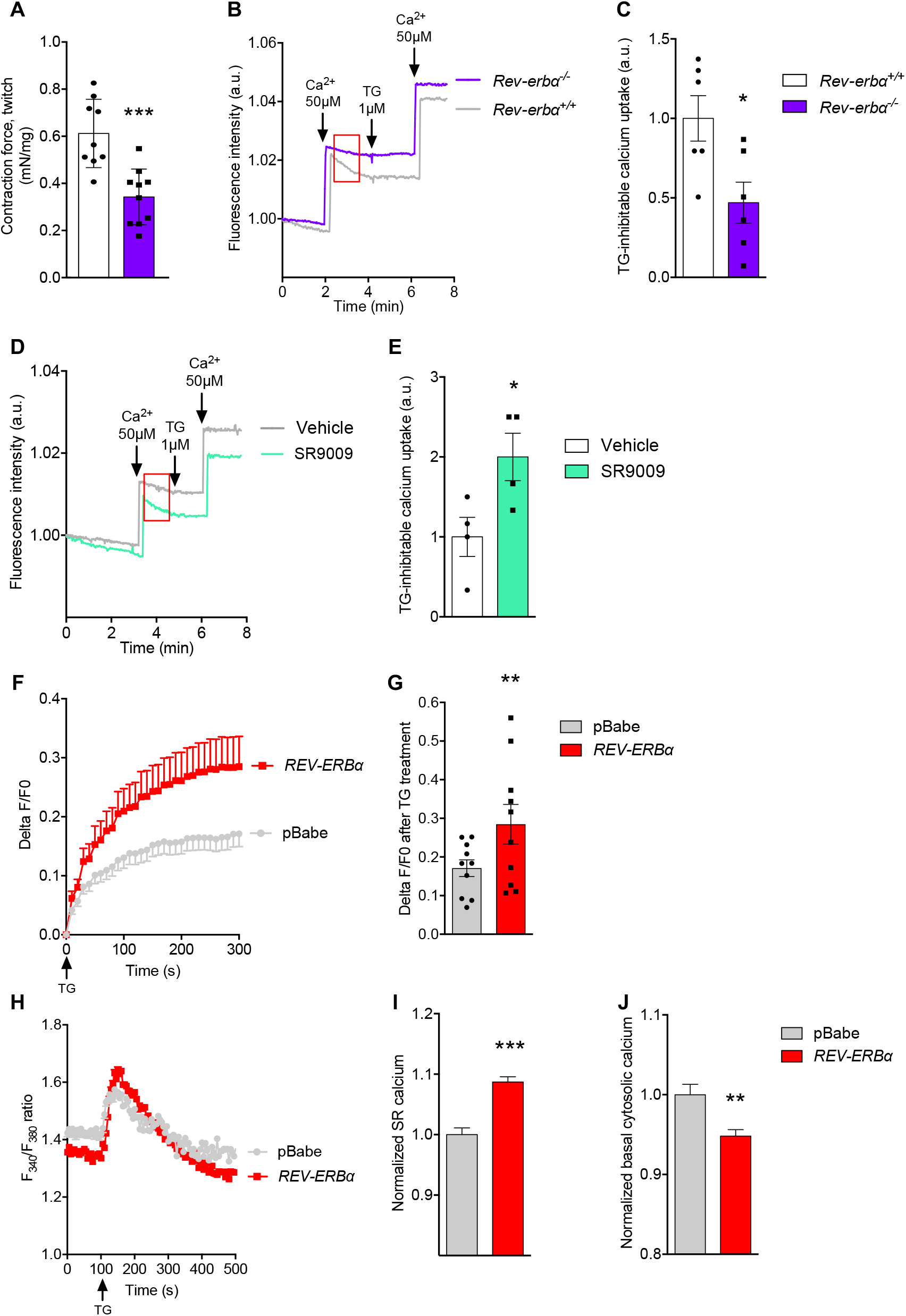
Rev-erb-α regulates SR Ca^2+^ homeostasis in skeletal muscle. (**A**) *In situ* measurement of *gastrocnemius-developed* force upon an electrical stimulus in wild-type (*Rev-erbα^+/+^*) and *Rev-erbα^-/-^* mice, ***p=0.0004 *vs. Rev-erbα^+/+^*, n=9-10, unpaired t-test. (**B**) Representative curves of SERCA-inhibitable Ca^2+^ uptake in microsomal fractions prepared from muscle from *Rev-erbα^+/+^* and *Rev-erbα^-/-^* mice. Decrease in fluorescence over time indicates Ca^2+^ uptake by the microsomal fraction. Arrows indicate Ca^2+^ solution (50μM) or thapsigargin (TG, 1μM) injections. Red rectangle indicates the region used for the slope calculation of fluorescence decrease. (**C**) Slopes of the decreasing fluorescence over time, indicative of the specific SERCA Ca^2+^ uptake in microsomal fraction obtained from *Rev-erbα^+/+^* and *Rev-erbα^-/-^* muscle. Data are represented as means ± SEM, n=6, *p=0.0203 *vs. Rev-erbα^+/+^*, unpaired t-test. (**D**) Representative curves of SERCA-inhibitable Ca^2+^ uptake in microsomal fraction prepared from muscle from vehicle-treated (vehicle) and SR9009-injected (SR9009) wild-type mice. (**E**) Slopes of the decreasing fluorescence over time. Data expressed as means ± SEM, n=4, *p=0.0408 *vs.* vehicle, unpaired t-test. (**F**) TG-induced Sarcoplasmic Reticulum (SR) Ca^2+^ release in control pBabe and *REV-ERBα* overexpressing C2C12 myotubes. Cells are loaded with Fluo4-AM to detect cytosolic Ca^2+^. SR Ca^2+^ content depletion is induced by TG (1μM). Results are expressed as means ± SEM of Delta F/F0 ratio, n=10. (**G**) Delta F/F0 ratio normalized to pBabe values, obtained 5min after TG-induced Ca^2+^ release and expressed as means ± SEM. n=10, **p=0.007 *vs.* pBabe, unpaired t-test. (**H**) Representative experiments of Fura-2/AM fluorescence intensity (ratio F340/F380) (Delta F/F0) over time of pBabe and *REV-ERBα* overexpressing cells. Calcium release from the SR was induced by adding 1μM Thapsigargin (TG). (**I**) Normalized SR calcium concentration (area under the curve), released upon TG treatment, in pBabe and REV-ERBα overexpressing cells. n>800 cells in each group, ***p<0.0001 *vs.* pBabe, unpaired t-test. (**J**) Normalized basal cytosolic calcium concentration (mean of the 100 first seconds) in pBabe and REV-ERBα overexpressing cells. n>800 cells in each group, **p=0.0013 *vs.* pBabe, unpaired t-test.

### Rev-erb-α controls Ca^2+^ homeostasis through direct repression of Myoregulin expression

We then aimed to identify the mechanism by which Rev-erb-α regulates calcium homeostasis. Because skeletal muscle SR Ca^2+^ homeostasis is mainly controlled by the Ryanodine Receptor RyR1, SERCA1 and SERCA2, we determined whether Rev-erb-α controls their expression. Whereas *Ryr1, Serca1* and *Serca2* expression was identical in *Rev-erbα^+/+^* and *Rev-erbα^-/-^* mice (Figures 2A-2C) as well as in pBabe and *REV-ERBα*-overexpressing cells (Figures 2D-2F), expression of *Mln*, a recently identified skeletal muscle-specific SERCA inhibitor (Anderson et al., 2015), was significantly higher in skeletal muscle from *Rev-erbα^-/-^* mice compared to control littermates (Figure 2G). In line, treatment with SR9009 *in vivo* to activate Rev-erb-α significantly decreased *Mln* expression (Figure 2H). Consistently, *REV-ERBα* overexpression or Rev-erb pharmacological activation with SR9009 decreased *Mln* expression in C2C12 cells (Figures 2I-2J), whereas cell treatment with the Rev-erb antagonist SR8278 induced *Mln* expression (Figure 2K).

**Figure 2.**
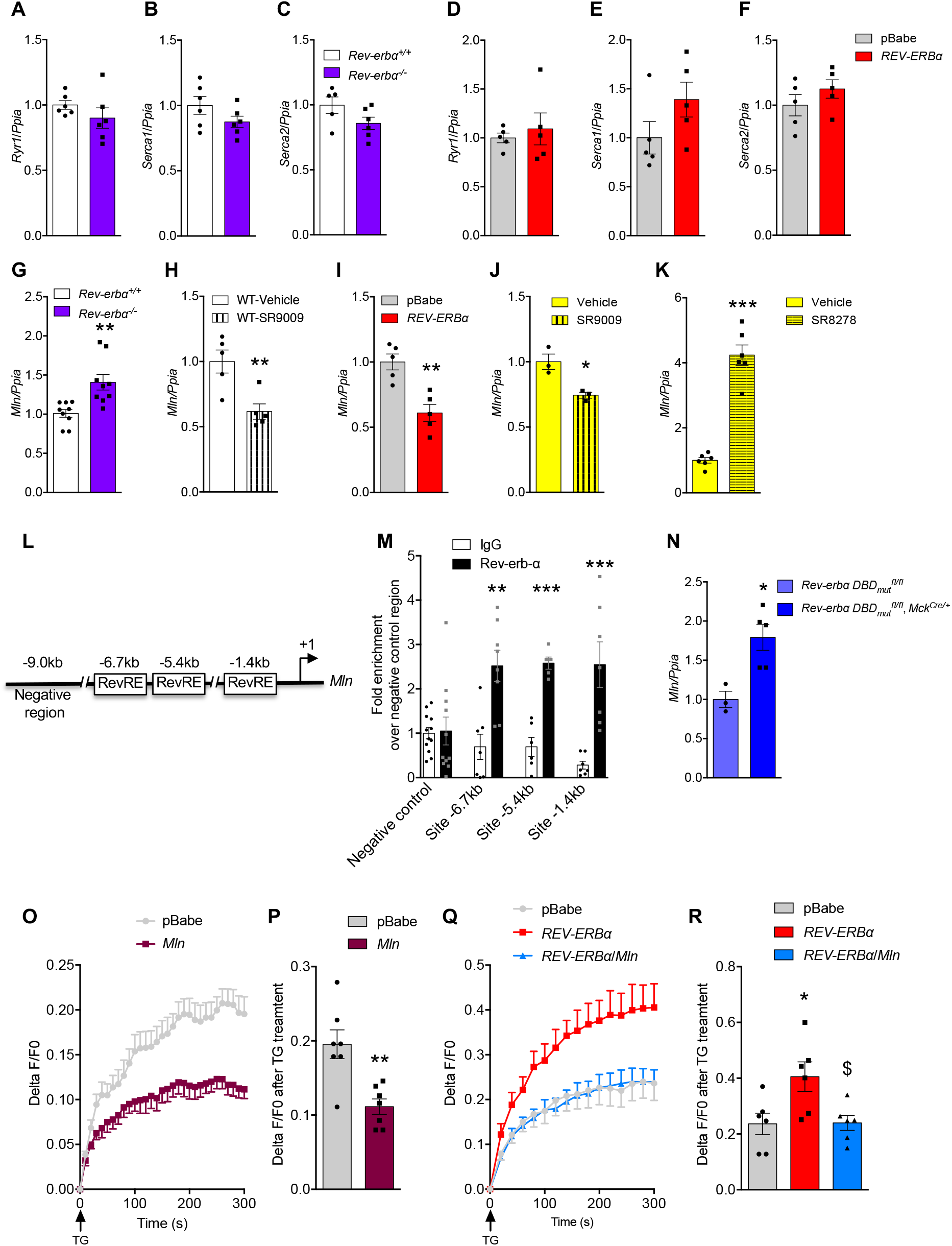
Rev-erb-α represses *Myoregulin (Mln*) expression through direct binding to its promoter. (**A**) *RyR1,* (**B**) *Serca1* and (**C**) *Serca2* gene expression in muscle from *Rev-erbα^+/+^* and *Rev-erbα^-/-^* mice (n=6, NS by unpaired t-test), (**D**) *RyR1,* (**E**) *Serca1* and (**F**) *Serca2* gene expression levels in pBabe and *REV-ERBα* overexpressing differentiated C2C12 (n=3, NS by unpaired t-test). *Mln* expression in (**G**) muscle from *Rev-erbα^+/+^* and *Rev-erbα^-/-^* mice (n=6, **p=0.0025 compared to *Rev-erbα^+/+^*, unpaired t-test), (**H**) muscle from SR9009 treated wild-type (WT) animals (n=5, **p=0.0069 compared to vehicle, unpaired t-test), (**I**) *REV-ERBα* overexpressing C2C12 (n=5, **p=0.0023 *vs.* pBabe, unpaired t-test), (**J-K**) C2C12 treated either with (**J**) 10μM of the Rev-erb agonist SR9009 (n=3, *p=0.0155) or (**K**) 10μM of the Rev-erb antagonist SR8278 (n=6, ***p<0.0001 *vs.* DMSO-treated cells, unpaired t-test). (**L**) Schematic representation of the *Mln* promoter indicating the presence of three putative Rev-erb-α Response Elements (RevRE), located ~1.4, ~5.4kb and ~6.7kb upstream the transcription initiation site. (**M**) Chromatin ImmunoPrecipitation analysis using an anti-Rev-erb-α antibody or control Immunoglobulin G (IgG) and specific primers targeting the three identified putative sites or a negative control region located ~9kb upstream the transcription initiation site. n=6-8 mice, data are means ± SEM, site −6.7kb **p=0.0017, site −5.4kb ***p<0.0001, site −1.4kb ***p=0.001 *vs.* IgG. (**N**) *Mln* expression in mice with muscle-specific expression of a mutated isoform of Rev-erb-α lacking the DNA binding domain (*Rev-erbα DBD_mut_^fl/fl^, MCK^Cre/+^*) and control *Rev-erbα DBD_mut_^fl/fl^* mice. n=3-5, *p=0.0139 *vs. Rev-erbα DBD_mut_^fl/fl^*, unpaired t-test. (**O**) Thapsigargin (TG)-induced Sarcoplasmic Reticulum (SR) Ca^2+^ release in pBabe and Mln overexpressing differentiated C2C12. Cells are loaded with Fluo4-AM and SR Ca^2+^ release is induced by the addition of 1μM TG. Results are expressed as means ± SEM of the Delta F/F0 ratio, n=7. (**P**) Peak fluorescence intensity of thapsigargin (TG)-induced Sarcoplasmic Reticulum (SR) Ca^2+^ release in pBabe *Mln* overexpressing differentiated C2C12, normalized to pBabe and expressed as mean ± SEM. n=7, **p=0.024 *vs.* pBabe, unpaired t-test. (**Q**) Thapsigargin (TG)-induced Sarcoplasmic Reticulum (SR) Ca^2+^ release in pBabe, *REV-ERBα* overexpressing and *REV-ERBα/Mln* overexpressing differentiated C2C12. Cells are loaded with Fluo4-AM and SR Ca^2+^ release is induced by the addition of 1μM TG. Results are expressed as means ± SEM of the Delta F/F0 ratio of 6 independent experiments. (**R**) Peak fluorescence intensity of thapsigargin (TG)-induced Sarcoplasmic Reticulum (SR) Ca^2+^ release in pBabe, *REV-ERBα* overexpressing and *REV-ERBα/Mln* overexpressing differentiated C2C12, normalized to pBabe and displayed as means ± SEM. n=6, *p<0.026 *vs.* pBabe, $p<0.0293 *vs.* REV-ERBα, 1-way ANOVA, Tukey’s multiple comparison test.

*In silico* analysis identified at least three putative Rev-erb Response Elements (RevRE) located at 1.4, 5.4 and 6.7kb upstream the *Mln* transcription start site (Figure 2L). Using Chromatin ImmunoPrecipitation (ChIP)-qPCR experiments performed on mouse skeletal muscle, we demonstrate that Rev-erb-α binds to these three regions (Figure 2M). To test whether direct Rev-erb-α binding to the *Mln* gene is required for its regulation, we used skeletal muscle-specific mutant mice expressing a DNA Binding Domain (DBD)-deficient Rev-erb-α protein. As observed in the *Rev-erbα^-/-^* mice, *Mln* expression was higher in skeletal muscle-specific mutant mice compared to wild-type floxed littermates (Figure 2N). Altogether, these data reveal that Rev-erb-α represses *Mln* gene expression by direct binding to its promoter.

To functionally demonstrate that MLN is key in the regulation of muscle Ca^2+^ homeostasis by Rev-erb-α, we aimed to restore MLN expression in *REV-ERBα* overexpressing cells. As expected, overexpression of *MLN* alone, by viral vector transduction in C2C12 myotubes, reduced SR Ca^2+^ stores compared to control pBabe cells (Figures 2O and 2P). More importantly, MLN overexpression in *REV-ERBα*-overexpressing cells normalized Ca^2+^ handling (Figures 2Q and 2R).

### Pharmacological Rev-erb activation alleviates the dystrophic phenotype in Duchenne Myopathy

Calcium homeostasis is known to be impaired in several myopathies (Rivet-Bastide et al., 1993; Vallejo-Illarramendi et al., 2014)(Vallejo-Illarramendi et al., 2014). To determine whether this could be due, at least in part, to a deregulation of Rev-erb-α and its downstream targets, we first analyzed its expression in publicly available microarray datasets from dystrophic muscles. Interestingly, we found that REV-ERB-α is expressed, albeit to significantly lower levels, in Duchenne Muscular Dystrophy (DMD) patients from different cohorts (Figure 3A, Supplemental Figures S1A-B). The same trend was observed in muscle biopsies from DMD patients kindly provided by the French Myobank (Supplemental Figure S1C). In line, *REV-ERBα* expression was reduced by ~30% in DMD muscle cells compared to human control myoblasts (Figure 3B). These data indicate altered Rev-erb-α expression, hence action, may be associated to the Duchenne dystrophy, pointing to Rev-erbα an interesting target specifically in this myopathy.

**Figure 3.**
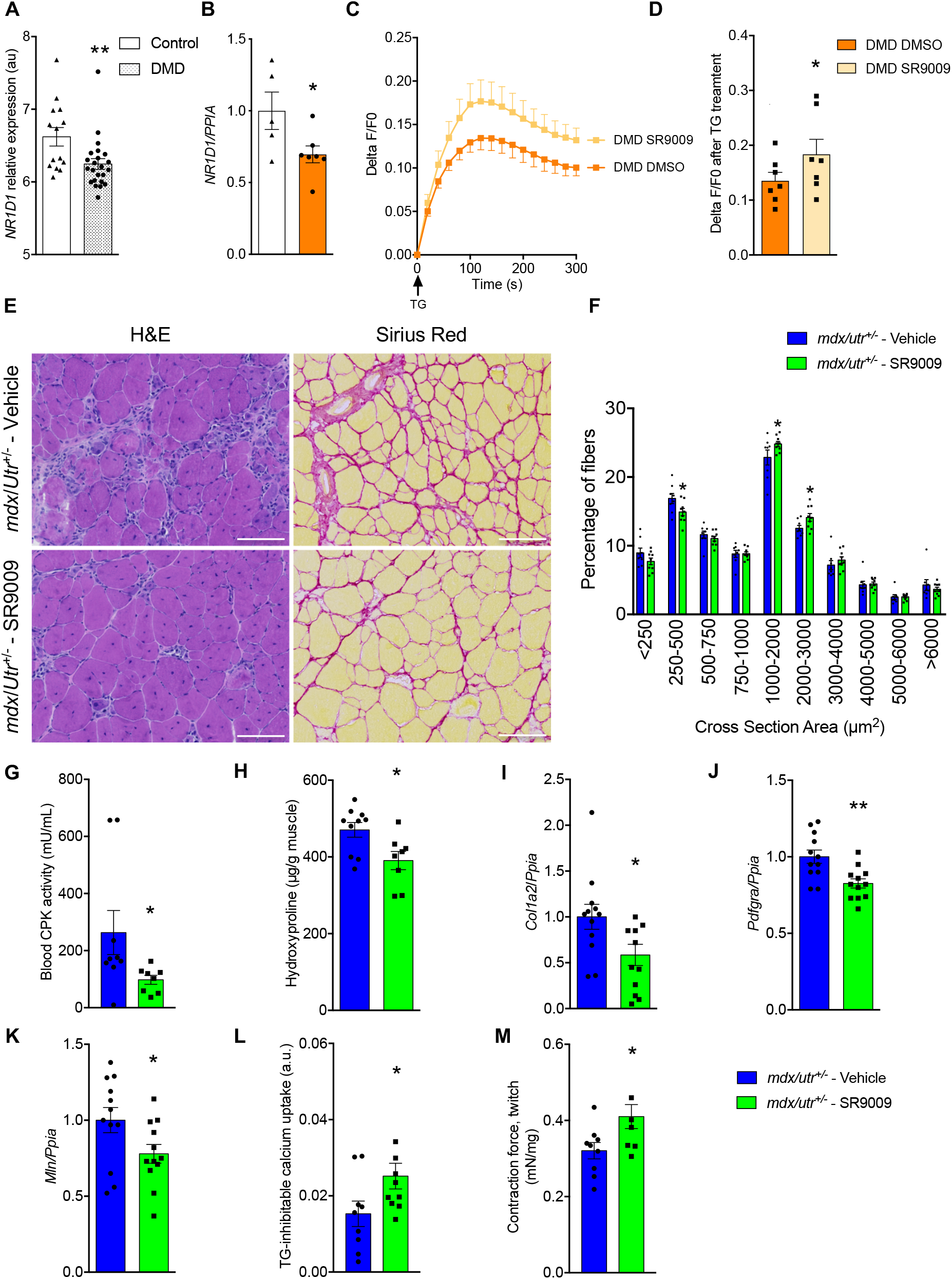
Rev-erb activation alleviates Duchenne Muscular Dystrophy features both in mice and human myoblasts. (**A**) *NR1D1* (*REV-ERB-α*) expression in muscle biopsies from controls (n=14) and patients suffering from Duchenne Muscular Dystrophy (DMD, n=23), **p=0.0092, unpaired t-test, data from GEO DataSets GSE6011. (**B**) *NR1D1* expression in control or DMD myoblasts, n=5-7. *p=0.0404 *vs.* control cells in panel B, unpaired t-test. (**C**) Representative curves and (**D**) peak fluorescence intensity of thapsigargin (TG)-induced Sarcoplasmic Reticulum (SR) Ca^2+^ release in myoblasts from controls or patients suffering from Duchenne Muscular Dystrophy (DMD) treated with SR9009 (10μM) or vehicle. Cells are loaded with Fluo4-AM and SR Ca^2+^ release is induced by the addition of 1μM TG. Results are expressed as means ± SEM of the Delta F/F0 ratio, n=3 controls, n=7 in both DMD groups. **p=0.0049 *vs.* control cells, unpaired t-test, $p=0.0138 *vs.* DMD DMSO, paired t-test. (**E**) Hematoxylin and eosin and Sirius red staining of *tibialis anterior* muscles obtained from vehicle- and SR9009-injected *mdx/Utr^+/-^* mice. Scale bars represent 100μm. (**F**) Myofiber cross-sectional area distribution (n=7-9), *p<0.05 vs. vehicle-treated *mdx/Utr^+/-^* mice. (**G**) Circulating Creatine PhosphoKinase (CPK) activity, n=8-9, *p=0.0329 *vs.* vehicle-treated *mdx/Utr^+/-^* animals. (**H**) Muscular hydroxyproline, n=8-10, *p=0.0172. **(I)** *Col1a2,* (**J**) *Pdgfra* and (**K**)*Mln* gene expression; n=8-12, *p=0.0314, *p=0.0164 and *p=0.0402, respectively. (**L**) SERCA activity (n=8-10) in muscular microsomes from *mdx/Utr^+/-^* mice treated for 20 days with SR9009 (100mg/kg) or vehicle; *p=0.0266. (**M**) *In situ* measurement of *gastrocnemius*-developed force (n=4-9, *p=0.0301).

Next, we thought to determine whether pharmacological Rev-erb-α activation might alleviate the dystrophic phenotype of DMD. We tested whether pharmacological REV-ERB activation by SR9009 may improve Ca^2+^ homeostasis in cells from patients suffering from DMD. A significantly higher SR Ca^2+^ release triggered by TG was measured in SR9009-treated compared to vehicle-treated DMD cells (Figure 3C-D).

To further assess whether pharmacological Rev-erb activation could improve muscle function in a pathological DMD model *in vivo, mdx/Utr^+/-^* mice, which closely recapitulate the features of the human disease, were daily injected with SR9009 for 20 days. Histological analysis revealed that tissue architecture was improved in SR9009-treated mice (Figure 3E and Supplemental Figure S2A), along with a mild, but significant, decrease in small fibers and an increase in medium/large fibers compared to vehicle-injected *mdx/Utr^+/-^* mice (Figure 3F). Circulating blood Creatine Phospho Kinase (CPK), which is a marker of muscle damage, was strongly decreased in the SR9009-treated group compared to vehicle-treated *mdx/Utr^+/-^* mice (Figure 3G). Several fibrosis markers, including Sirius red staining (Figure 3E), muscle hydroxyproline quantity (Figure 3H), *Colla2* (Figure 3I) and *Pdgfra* (Figure 3J) expression, were also reduced by SR9009 treatment. Next, we evaluated whether SR9009 treatment was able to improve muscle calcium homeostasis and function in this model of myopathy. As expected based on the data from the genetic models of deletion or over-expression of Rev-erb-α specifically in skeletal muscle, daily injection of SR9009 for 20 days reduced *Mln* expression in muscle from *mdx/Utr^+/-^* mice (Figure 3K), whereas SERCA activity was improved (Figure 3L), both being strongly correlated (Supplemental Figure S2B). More importantly, gastrocnemius muscle-developed contraction force was significantly ameliorated by SR9009 treatment in *mdx/Utr^+/-^* mice compared to vehicle-injected littermates (Figure 3M).

In conclusion, a 20-day Rev-erb agonist treatment improved SR homeostasis and muscle function in a mouse model of Duchenne myopathy.

## Discussion

Our data demonstrate that Rev-erb-α improves calcium homeostasis in skeletal muscle by directly controlling *Mln* expression, hence SERCA activity. We also report that Rev-erb-α pharmacological activation by synthetic ligands can reveal therapeutic interest since it improves calcium homeostasis in human cells from DMD patients and alleviates the myopathy phenotype in *mdx/Utr^+/-^* mice.

RyR1 and SERCA1 are the two major SR proteins controlling Ca^2+^ fluxes in skeletal muscle. Nevertheless, neither RyR1 nor SERCA1 expression was impacted by Rev-erb-α. Therefore, we focused on Mln, the main glycolytic/mixed muscle endogenous SERCA inhibitor (Anderson et al., 2016). Mln is a recently discovered 46-amino acid micropeptide that forms a single transmembrane alpha helix and interacts with the skeletal muscle SERCA1 isoform to inhibit its pumping activity, thereby decreasing SR Ca^2+^ content (Anderson et al., 2015). Here, we identified Rev-erb-α as a new direct transcriptional repressor of Mln gene expression. Indeed, we demonstrated that Rev-erb-α binds to three RevRE located in the *Mln* promoter *via* a functional DNA binding domain. By modulating Ca^2+^ handling, Mln was proposed to modulate skeletal muscle contractile activity and to represent a promising drug target for improving Ca^2+^-related skeletal muscle disorders and muscle performance (Anderson et al., 2015). Consistently, *Mln* deletion in mice improves skeletal muscle performance (Anderson et al., 2015). Yet, modulators of *Mln* expression remained to be identified. Interestingly, pharmacological Rev-erb activation, which we have shown in previous studies to improve muscle performance in non-pathological contexts (Woldt et al., 2013) and to block glucocorticoid-induced muscle wasting (Mayeuf-Louchart et al., 2017), is able to repress *Mln* expression. Therefore, we bring novel insights into the molecular mechanisms by which Rev-erb-α exerts beneficial effects on muscle function and uncover a novel pathway to control *Mln*, hence skeletal muscle Ca^2+^ handling and likely contractile function.

We have demonstrated that *REV-ERB-α* is expressed in DMD cells, albeit to a lower extent compared to control human myotubes, suggesting that increasing Rev-erb activity could represent a new therapeutic option in myopathies. In muscle of patients suffering from DMD, the absence of dystrophin causes muscular contraction impairment with altered Ca^2+^ handling, *i.e.* raised cytosolic Ca^2+^ concentrations and depletion of SR Ca^2+^ stores due to impaired uptake capacity (Vallejo-Illarramendi et al., 2014; Voit et al., 2017). Here, we confirm these data and we further demonstrate that pharmacological activation of Rev-erb by a synthetic ligand improves SR Ca^2+^ content in myoblast cells obtained from DMD patients. This was also observed in vivo in a DMD mouse muscle in which pharmacological activation of Rev-erb significantly improved muscle histology and reduced damage markers and fibrosis. By itself, and consistent with other studies showing that improving Ca^2+^ homeostasis mitigates DMD (Mázala et al., 2015; Voit et al., 2017), reduction of MLN expression by pharmacological Rev-erb activation may contribute to the improved muscle contractility observed in myopathic mice.

Others reported that Rev-erb antagonism with SR8278 may also improve muscle function, reduce fibrosis and increase mitochondrial biogenesis in *mdx* mice (Welch et al., 2017). This study is in apparent contradiction with the present results and with results that, we and these authors, have previously published demonstrating that Rev-erb agonism with SR9009 improves muscle mitochondrial function and exercise capacity in healthy mice (Woldt et al., 2013). In addition, we and others have reported that Rev-erb-α positively controls skeletal muscle mass by counteracting both autophagy (Woldt et al., 2013) and proteasomal-associated fiber atrophy (Mayeuf-Louchart et al., 2017) and by promoting myoblast differentiation through mTORC1 signaling pathway activation (Maayan et al., 2020), again supporting a positive action of Rev-erb-α in skeletal muscle. Compensatory mechanisms may interfere as both Rev-erb-β overexpression and knock-down were reported to lead to a similar increase in mitochondrial biogenesis (Amador et al., 2018). Moreover, SR9009 as well as SR8278 target both Rev-erb-α and Rev-erb-β (Kojetin and Burris, 2014) and may also exert Rev-erb-independent activities (Dierickx et al., 2019). In the present study, we have used genetic models of deletion or over-expression of Rev-erbα specifically in skeletal muscle and in mouse and human myoblasts to support our model and validate the role of Rev-erb-α in ameliorating muscle calcium handling and improving dystrophy. While this is possibly one reason for the apparent discrepancy between our results and others (Welch et al., 2017), it should also be noted that we used a different mouse model of muscle dystrophy. While the *mdx* mouse is widely used, it is a very mild model far from the human Duchenne myopathy phenotype (Larcher et al., 2014). In contrast, we used the *utr^+/-^ mdx* model that presents a profound phenotype more relevant to the human situation, which may also explain why different results were obtained. We also showed, for the first time, that this pertains to human myoblasts from Duchenne patients. Moreover, although it would be impossible to disentangle the two pathways, the beneficial effects of Rev-erb-α on muscle function could also be related to the combination of two mechanisms. Indeed, by improving mitochondrial function (Woldt et al., 2013), Rev-erb activation could lead to higher ATP availability for calcium pumps and myofibrillar proteins. Even though, for the above-mentioned reasons, the current ligands cannot be used in the clinic, our results advocate for further development of more selective Rev-erb-α-activating drugs, which could be of interest in the treatment of myopathies, and likely other muscle disorders characterized by altered Ca^2+^ homeostasis.

## Materials and Methods

### Study design

In the primary objective of our study, genetically-engineered and pharmacologically-treated cells and mice were used to determine whether Rev-erb-α modulates calcium homeostasis in the SR. The translational impact of our finding was then tested in human muscle cells obtained from patients suffering from DMD and in a mouse model for Duchenne myopathy. Based on age and weight, animals were randomly assigned to the different experimental groups. The number of samples for *in vivo* and *in vitro* assays was based on our experience and publications in the field.

### Cell culture and treatments

C2C12 cells (ATCC, Manassas, Virginia, USA) were cultured in high glucose DMEM (41965039, Gibco, Thermo Fischer Scientific, Waltham, Massachusetts, USA) supplemented with 10% fetal bovine serum and 0.4% gentamycin and differentiated by replacing the previous medium with DMEM supplemented with 2% horse serum and 0.4% gentamycin for 5 days. Myoblasts from control and DMD patients, kindly given by Myobank-AFM (Myology Institute, Pitié-Salpêtrière Hospital, Paris, France) were cultured in DMEM supplemented with 20% fetal calf serum and 0.2% primocin.

Generation of REV-ERBα and myoregulin (Mln) overexpressing C2C12 was performed as previously described (Anderson et al., 2015; Woldt et al., 2013). Briefly, mouse Mln and human REV-ERBα coding sequences were inserted into the pBabe plasmid (Addgene, Cambridge, Massachusetts, USA) by using BamHI-SalII restriction sites. REV-ERBα and Mln or empty pBabe plasmids were transfected into Phoenix cells using JetPEI (Polyplus, Illkirch-Graffenstaden, France). Next, the supernatant of Phoenix cell culture was incubated with C2C12 cells, leading to their infection by retroviruses. The selection was done by a 15-day treatment with puromycin for REV-ERBα overexpressing cells, and neomycin for Mln overexpressing cells.

Pharmacological modulation of Rev-erb was obtained by adding in the culture medium either the synthetic agonist SR9009 (10μM) or the synthetic antagonist SR8278 (10μM). TG was dissolved in DMSO, which was added at the same concentration in control conditions.

### Mice housing and treatments

All mice were housed in our animal facility with a 12h/12h light/dark cycle and had free access to food and water. *Rev-erbα*-deficient mice (*Rev-erbα^-/-^*) and skeletal muscle-specific *Rev-erbα* DBD mutant mice (*Rev-erbα DBDmut^fl/fl^, MCK^Cre/+^*) expressing a truncated Rev-erb*α* lacking the DBD were generated as previously described (Woldt et al., 2013; Zhang et al., 2015) and compared to respective control littermates.

*Gastrocnemius* muscles were collected and were either flash-frozen in liquid nitrogen or rapidly frozen using isopentane cooled with liquid nitrogen for immunostaining, or freshly processed for microsome preparation. The effect of pharmacological Rev-erb activation on SR calcium uptake was tested in gastrocnemius muscle collected from wild-type mice treated with SR9009 (100 mpk) or its vehicle twice daily for 3 days (Mayeuf-Louchart et al., 2017).

To evaluate the therapeutic potential of Rev-erb activation in Duchenne myopathy, twenty five-week old *mdx/Utr^+/-^* (McDonald et al., 2015) mice were treated with SR9009 (100mg/kg, once a day for 20 days) or vehicle. All the described procedures were approved by the local ethics committee (CEEA75).

### Creatine PhosphoKinase activity

Blood CPK activity was measured with the creatine kinase assay kit (ab155901, Abcam, Cambridge, United Kingdom), according to manufacturer’s instructions.

### Muscular hydroxyproline assay

4-hydroxyproline, a major component of collagen, was detected by the use the assay kit MAK008 (Sigma-Aldrich, St. Louis, Missouri, USA), according to manufacturer’s instructions. Briefly, muscle (10 mg) was homogenized in 100 μL of water and hydrolysis was started by adding 100 μL of 12 M HCl. After 3 hours at 120°C, samples were spun down at 10,000 *g*. 20 μL of the resulting supernatant were transferred in a 96-well plate and evaporated under vacuum. 100 μL of chloramine T/oxidation buffer mixture was added into the wells. Then, 100 μL of DMAB reagent were added. Plate was incubated for 90 minutes at 60°C. Absorbance was measured at 560nm and compared to hydroxyproline standards.

### In Situ Contractile Properties of the Gastrocnemius Muscle

Mice were deeply anesthetized with intraperitoneal injections of ketamine (50 mg.kg^-1^) and dexmedetomidine (Domitor, 0.25 mg.kg^-1^). The dissection protocol was previously described (Picquet and Falempin, 2003). Briefly, all the muscles of the right hindlimb were denervated, except the gastrocnemius muscle, which was isolated from surrounding tissues. Then, the limb was immersed in a bath of paraffin oil thermostatically controlled (37°C), and fixed with bars and pins. The gastrocnemius muscle was maintained in a horizontal position and its distal tendon was connected to a force transducer (Grass FT 10, Grass Instruments, West Warwick, Rhode Island, USA). The muscle length was adjusted to produce a maximal twitch peak tension (Pt). Contractions were induced by stimulation of the sciatic nerve (0.2ms pulses) through bipolar platinum electrodes at twice the minimum voltage required to obtain the maximal twitch response. At the end of the recording session, the muscle was removed for determination of muscle wet weight, frozen in liquid nitrogen and stored at −80°C.

### RT-qPCR analysis

ARN were extracted from mouse muscle, C2C12 and human myoblasts seeded in 6-well plates or from *gastrocnemius* muscle, according to the Trizol (Invitrogen, Thermo Fischer Scientific, Waltham, Massachusetts, USA)/Chloroform/Isopropanol protocol. After DNase treatment, cDNA was obtained using the High-Capacity cDNA Reverse Transcription Kit (Life Technologies, Carlsbad, California, USA). qPCRs were realized using SYBR® Green Real-Time PCR Master Mix kit (Agilent Technologies, Santa Clara, California, USA) and a MX3005 apparatus (Agilent Technologies, Santa Clara, California, USA). Mouse and human-specific primers are recapitulated in supplemental tables S1 and S2, respectively. Gene expression was normalized to cyclophilin A (*Ppia*).

### SERCA-dependent Ca^2+^ uptake

*Gastrocnemius* muscles were collected and homogenized at 4°C in a dedicated buffer (Tris-HCl pH7 1M, sucrose 8%, PMSF 1mM, DTT 2mM) with a Polytron (Kinematica AG, Malters Switzerland). Samples were then centrifuged at 1,300*g* at 4°C for 10min in order to remove nuclei. The supernatant obtained after a second centrifugation (20,000*g*, 4°C, 20min) corresponds to the enriched microsomal fraction. 150μg proteins were placed in calcium uptake buffer (CaCl_2_ 120μM, EGTA 150μM, Tris-HCl 30mM pH7, KCl 100mM, NaN_3_ 5mM, MgCl_2_ 6mM, oxalate 10mM) and put in 2mL-chambers of the Oxygraph-2k (Oroboros Instruments, Innsbruck, Austria) equipped with the fluorescence LED2-module. Calcium green probe (1μM) and ATP (5mM) were added and fluorescence was measured over time (λex 506nm, λem 531nm). Ca^2+^ (50μM) pulse was then injected into the chambers. Finally, TG (1μM) was added in order to ensure that SERCA-dependent Ca^2+^ uptake was measured. We calculated the slope of the fluorescence intensity decrease subtracted with the residual slope measured in the presence of TG reflects SERCA-dependent Ca^2+^ uptake.

### SR Ca^2+^ content in C2C12 cells

Experiments were conducted following a technical protocol adapted from Ducastel *et al.* (Ducastel et al., 2020). C2C12 cells and human myoblasts were plated in a 96-well plate (20 000/well) and differentiated for 4 days. Then, medium was replaced for 24 hours by low Ca^2+^ concentration Locke’s Buffer (NaCl 154mM, NaHCO_3_ 4mM, KCl 5mM, CaCl_2_ 2H_2_O 0.1mM, MgCl_2_ 6H_2_O 1mM, Glucose 5mM, Hepes 10mM, pH7.4), as previously described (Brandman et al., 2007). To detect cytosolic Ca^2+^, myotubes were then loaded with Fluo4-AM (λex 490nm, λem 516nm) for 30min at 37°C, with 5% CO_2_ in free Ca^2+^ Locke’s buffer. Following 2 washes with 2.3mM Ca^2+^ Locke’s buffer, TG (1μM) was added in order to deplete SR Ca^2+^ store. Fluorescence intensity was immediately recorded every 10 seconds during 5min using a microplate reader (Infinite 200 pro, Tecan, Männedorf, Switzerland) in order to estimate SR Ca^2+^ content until stabilization.

### Calcium imaging

Cells were grown on glass bottom dishes to carry out calcium imaging experiments. Ratiometric dye Fura-2/AM (F1221, Invitrogen, Thermo Fischer Scientific, Waltham, Massachusetts, USA) was used as a Ca^2+^ indicator. Cells were loaded with 2μM Fura-2/AM for 45 min at 37°C and 5% CO_2_ in corresponding medium and subsequently washed three times with external solution containing (in mM): 140 NaCl, 5KCl, 1 MgCl_2_, 2 CaCl_2_, 5 Glucose, 10 Hepes (pH 7.4). The glass bottom dish was then transferred in a perfusion chamber on the stage of Nikon Eclipse Ti microscope (Nikon, Minato City, Tokyo, Japan). Fluorescence was alternatively excited at 340 and 380 nm with a monochromator (Polychrome IV, TILL Photonics GmbH, Kaufbeuren, Germany) and captured at 510 nm by a QImaging CCD camera (QImaging, Teledyne Photometrics, Tucson, Arizona, USA). Acquisition and analysis were performed with the MetaFluor 7.7.5.0 software (Molecular Devices Corp., San Jose, California, USA).

### Tissue histology

Cross Sectional Area was analyzed as previously described (Mayeuf-Louchart et al., 2018, 2017). Conventional hematoxylin-eosin (HE) staining was performed to describe histological status of muscle sections (Hardy et al., 2016). Sirius red staining was performed to describe fibrosis (Forand et al., 2020).

### ChIP experiment

ChIP assays were performed as previously described (Pourcet et al., 2018) with minor modifications as follows. *Gastrocnemius* muscles from wild-type C57/Bl6 mice were homogenized in LB1 buffer (Hepes-KOH 10mM pH7.5, NP-40 0.5%, MgCl_2_ 5mM, DTT 500μM, Cytochalasin B 3μg/mL, protease inhibitor cocktail) and cross-linked with 1% paraformaldehyde for 10min at room temperature. Chromatin was sheared during 90min using the Bioruptor (Diagenode, Liège, Belgium) coupled to a watercooling system and subsequently concentrated with centricon 10kDa column (Millipore, Burlington, Massachusetts, USA). 50μg of chromatin were immunoprecipitated overnight at 4°C with an antibody against Rev-erb-α (13418S, Cell signaling Technology, Danvers, Massachusetts, United States). BSA/yeast tRNA-blocked Protein A/G dynabeads (Invitrogen, Thermo Fischer Scientific, Waltham, Massachusetts, USA) were then added for 6h at 4°C while agitating and washed. Cross-linking was reversed by incubating precipitated chromatin overnight at 65°C. DNA was purified using the QIAquick PCR purification kit (Qiagen, Hilden, Germany) and was analyzed by qPCR using the Brilliant II SYBR Green QPCR Master Mix (Agilent Technologies, Santa Clara, California, USA) and specific primers (Supplemental Table S3).

### Statistical analyses

The number of sampled units, n, is reported in each figure legend. Values are means ± sem. The analysis was performed with GraphPad Prism software 5.0. One-way ANOVA followed by Tukey post-hoc tests are carried out in order to establish statistical significance when comparing three groups or more. The influence of Rev-erb-α expression was tested by two-way ANOVA followed by Sidak’s multiple comparisons test. Unpaired or paired Student t-tests were used to compare two groups, as indicated in figure legends. Significant effects are indicated as follows p<0.05 (*), p<0.01 (**), p<0.001 (***).

## Data availability

The publicly available GSE data were analyzed by the GEO2R tool available on the NCBI website (https://www.ncbi.nlm.nih.gov/geo/geo2r/). Benjamini & Hochberg (False discovery rate) was applied to the p-values. Data were then analyzed on GraphPad Prism 9.0.

## Author contributions

AB, CD and SL conceived and designed the experiments, AB, CD, AML, BP, YS, KK, AH, MG, CG, VM, SD, MC, QT, MZ, JB, AF and LF acquired and analyzed experiments, AB, CD, AML, BP, YS, KK, NP, FPR, BB, HD and SL interpreted data, AB, HD and SL wrote the original draft manuscript, AB, CD, BP, AML, YS, KK, NP, FPR, BB, BS, HD and SL reviewed and edited the manuscript, all authors approved the final version.

## Acknowledgments

The authors thank Myobank-AFM-Myology Institute (BB-0033-00012) for providing human myoblasts. A.B. was supported by a PhD scholarship from Lille University-Région Hauts-de-France and by EGID funds. Q.T. is supported by a PhD scholarship from Inserm-Région Hauts-de-France. M.Z. was supported by a PhD scholarship from Fondation pour la Recherche Médicale FRM (FDT20170739031). AML is supported by AFM (AFM N°22281). The authors acknowledge funding supports from INSERM, Contrat Plan Etat Région (CPER), Région Hauts-de-France/FEDER, CTRL-Lille Pasteur Institute, the European Genomic Institute for Diabetes (E.G.I.D., ANR-10-LABX-46), the European Foundation for the Study of Diabetes (EFSD), the Fondation Francophone pour la Recherche sur le Diabète (FFRD), sponsored by Fédération Française des Diabétiques (AFD), AstraZeneca, Eli Lilly, Merck Sharp & Dohme (MSD), Novo Nordisk & Sanofi. BS is a recipient of an Advanced ERC Grant (694717).

---

**Supplemental Figure S1.**
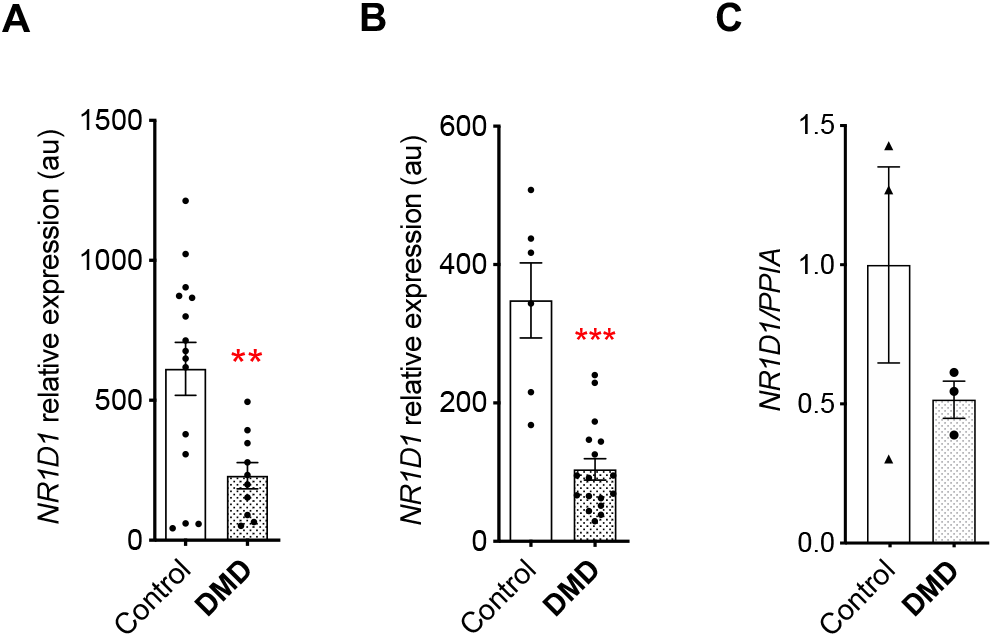
*REV-ERBα* (*NR1D1*) expression in muscles from patients suffering from Duchenne Muscular Dystrophy (DMD). Data from (**A**) GSE3307 probe 204769, (**B**) GSE109178 probe 31637. **p<0.01, ***p<0.001 *vs*. control, unpaired t-test. (**C**) RTqPCR results obtained in dorsal muscles from control and DMD patients provided by the French Myobank, n=3 samples in each group.

**Supplemental Figure S2.**
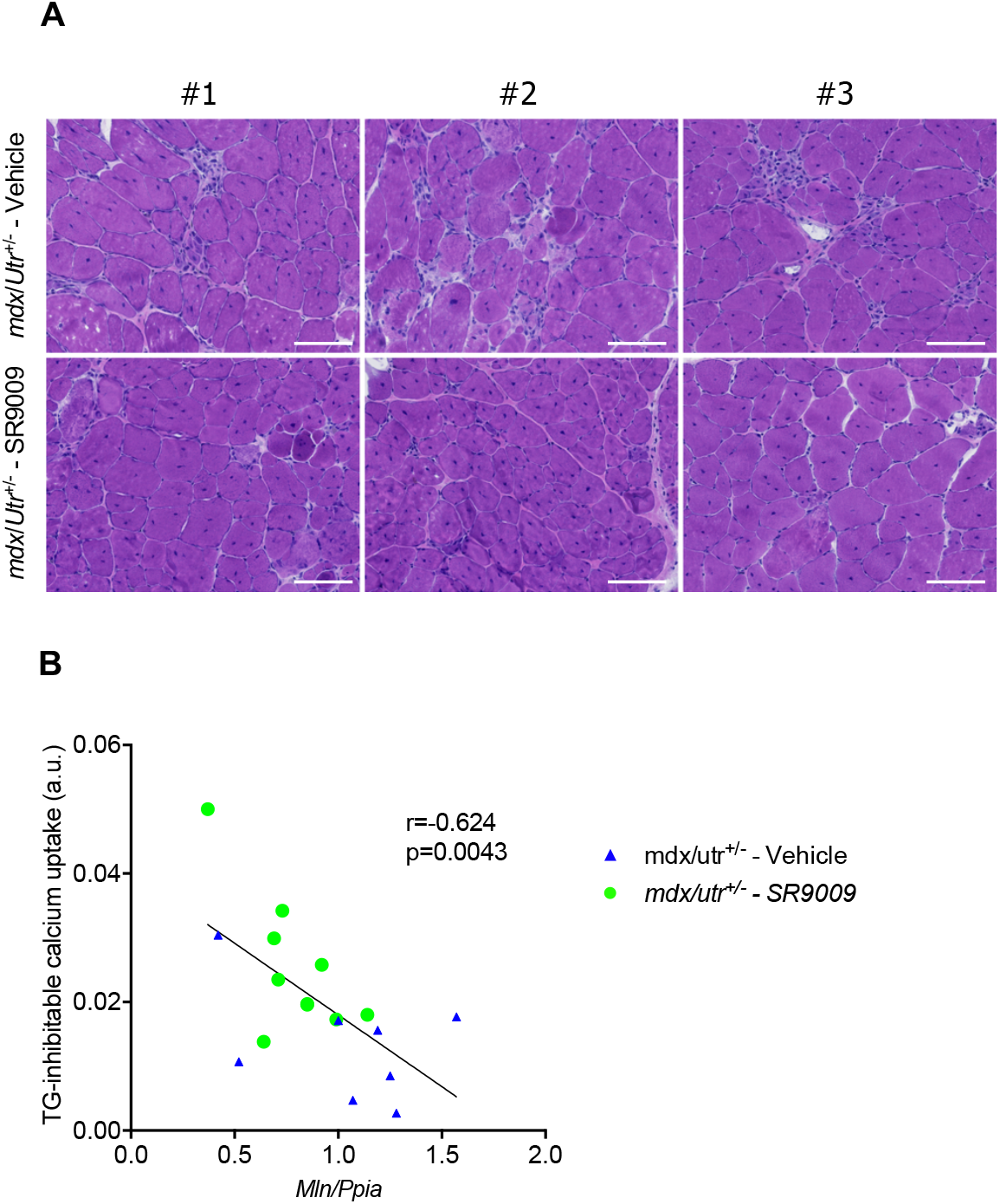
Effects of SR9009 on muscles from *mdx/Utr^+/-^* mice. (**A**) H&E staining on *tibialis anterior* sections from three different (#1, #2, #3) *mdx/Utr^+/-^* mice treated with SR9009 or vehicle for 20 days. Scale bars indicate 100μm. (**B**) Pearson correlation analysis between *Mln* expression and SERCA activity in muscle from vehicle- or SR9009-treated *mdx/Utr^+/-^* mice.

**Supplemental Table S1:**
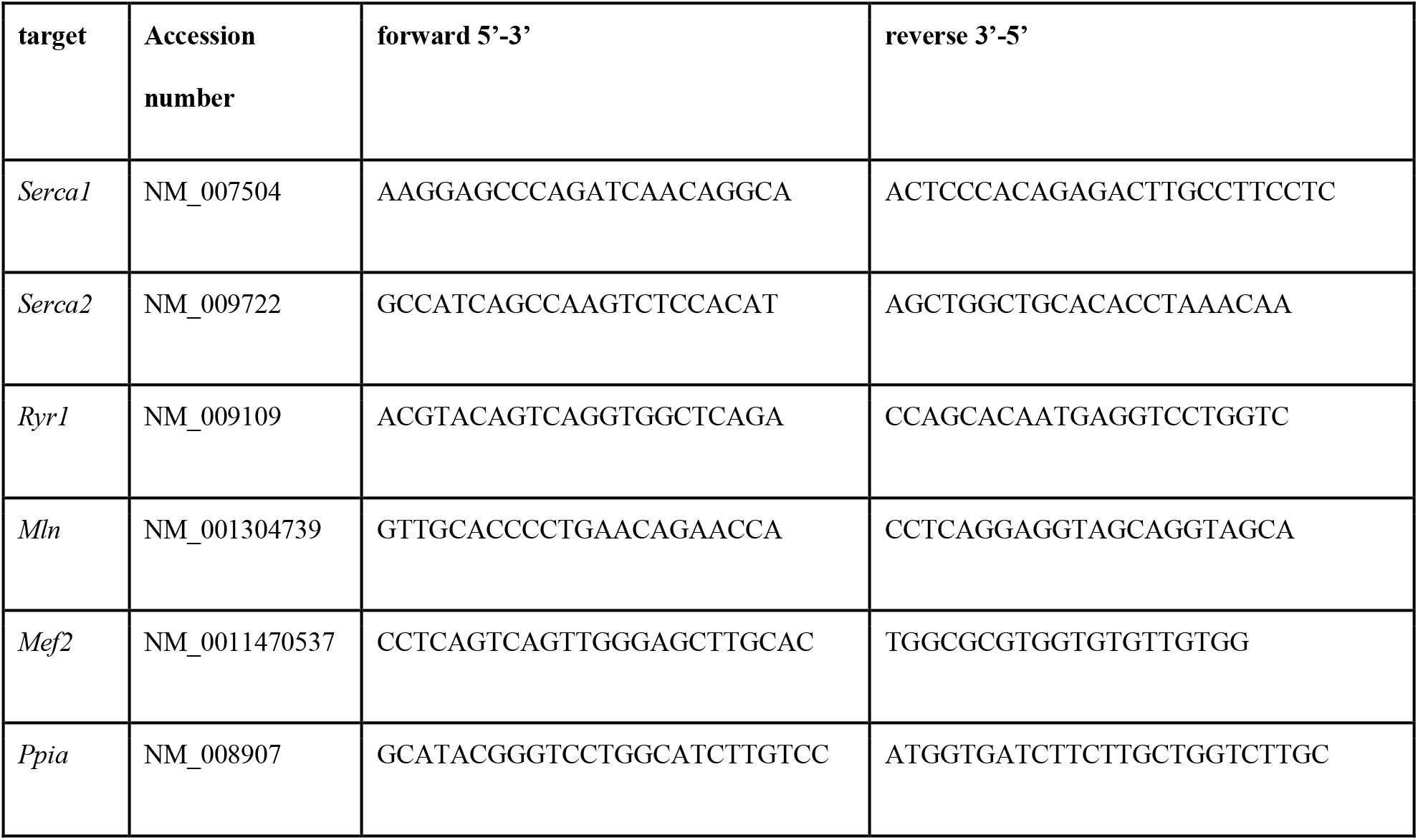
Mouse RTqPCR primers

**Supplemental Table S2:**
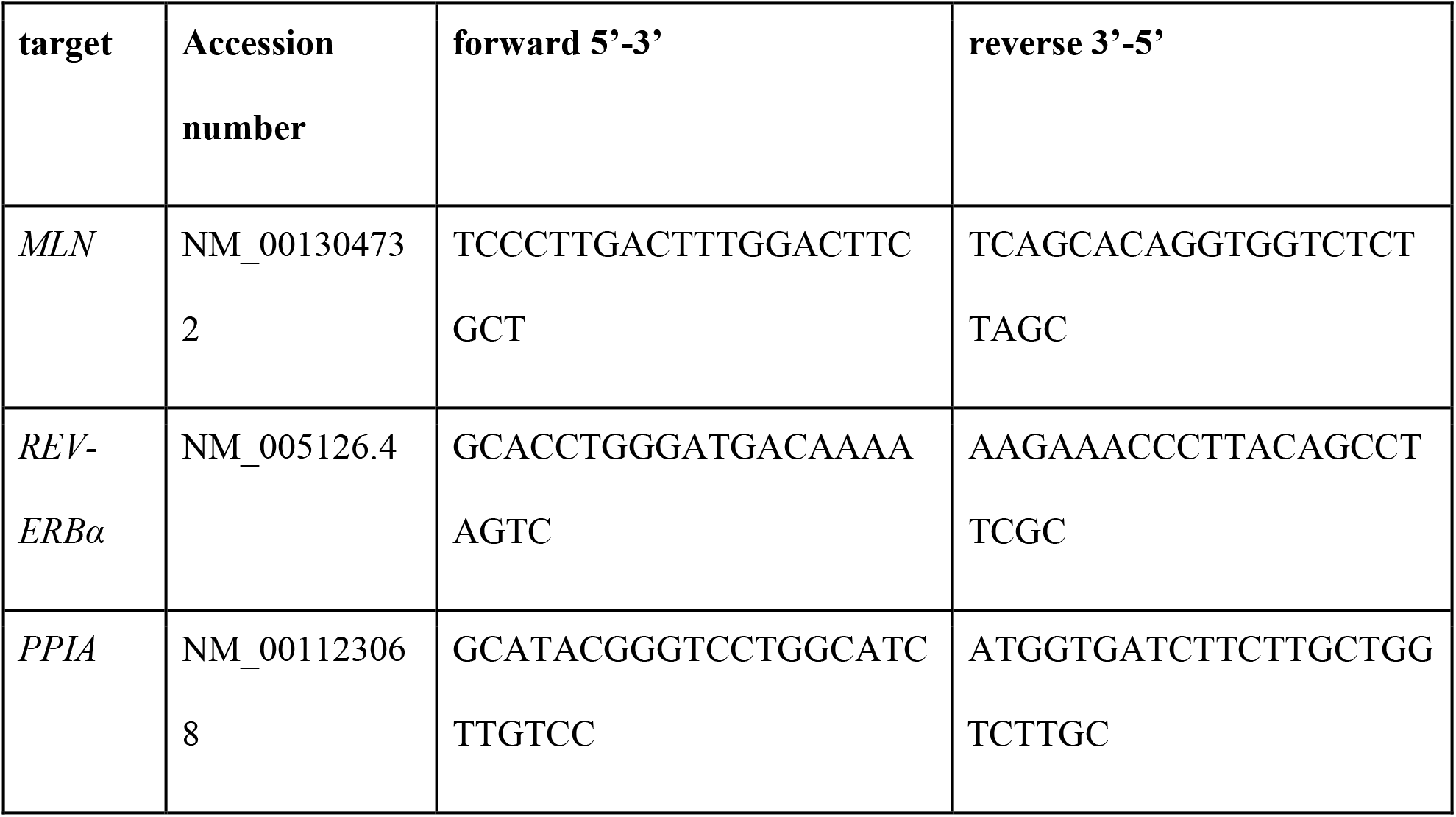
Human RTqPCR primers

**Supplemental Table S3:**
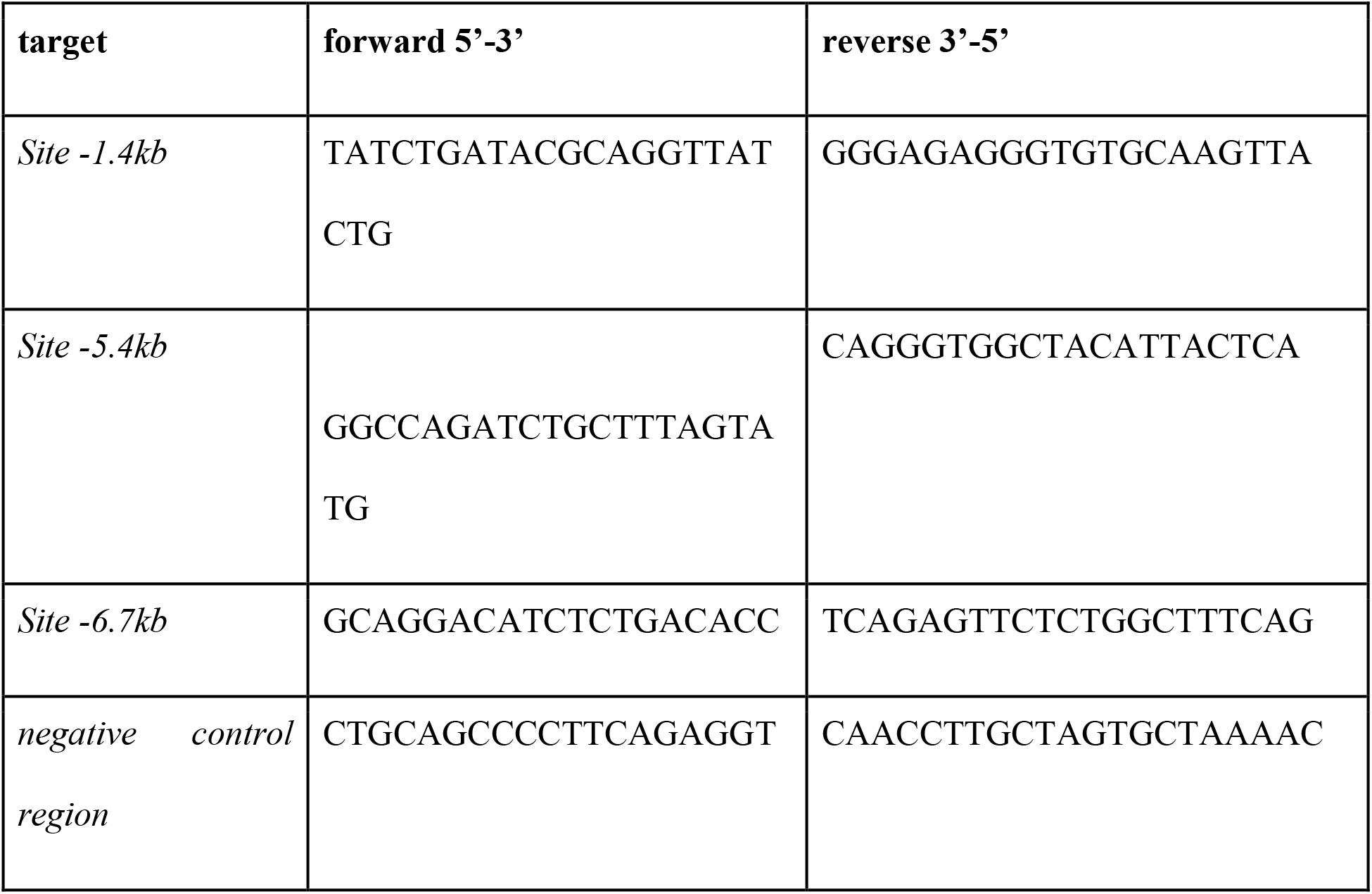
ChIP qPCR primers

